# Treatment of age-related visual impairment with a mitochondrial therapeutic

**DOI:** 10.1101/2020.11.06.371955

**Authors:** N.M. Alam, R.M. Douglas, G.T. Prusky

**Affiliations:** Burke Neurological Institute, 785 Mamaroneck Avenue, White Plains, New York 10605; University of British Columbia, Department of Ophthalmology and Visual Sciences, 2550 Willow Street, Vancouver, BC Canada V5Z 3N9; Weill Cornell Medicine, Department of Physiology and Biophysics, 1300 York Avenue, New York, NY 10065

## Abstract

Age-related visual decline and disease due to neural dysfunction are major sources of disability that have resisted effective treatment. In light of evidence that visual impairment and mitochondrial dysfunction advance with age, we characterized age-related decline of spatial visual function in mice, and investigated whether improving mitochondrial function could treat it. Impaired photopic acuity measured with a virtual opto-kinetic system emerged near 18 months, and declined to ∼40% below normal by 34 months. Daily application of the synthetic peptide SS-31, which has high selectivity for mitochondrial membranes that contain cardiolipin, and promotes efficient electron transfer, was able to mitigate visual decline from 18 months. Daily application from 24 months, when acuity was reduced by ∼16%, reversed visual decline and normalized function within 2 months; recovered function that persisted for at least 3 months after treatment was withdrawn. A single treatment at 24 months also delayed subsequent visual decline. Daily application from 32 months took longer to affect change, but enabled substantial improvement within 2 months. The effects of age and SS-31 treatment on contrast sensitivity was similar to those on acuity, systemic and eye drop applications of SS-31 had comparable effects, scotopic spatial visual function was largely unaffected by age or treatment, and altered function was independent of variation in optical clarity. These data indicate that SS-31 treatment adaptively alters the aging visual system, and provide a rationale to investigate whether mitochondrial dysfunction is a treatable pathophysiology of human visual aging and age-related visual disease.

**TRANSLATIONAL IMPACT:** *Clinical issue:* Age-related visual impairment is a major source of disability. Aging invariably leads to optical dysfunction related to inflexibility (presbyopia) or clouding (cataracts) of the lens, and neural dysfunction; each of which compromises the ability to resolve detail (acuity) and differences in luminance (contrast sensitivity) in visual scenes. Age is also a predisposition to develop blinding visual diseases that have a neurological origin, such as glaucoma, diabetic retinopathy, and age-related macular degeneration. Whereas, age-related optical problems can often be corrected with eyewear or surgical lens replacement, we lack sufficient understanding of the natural course of visual aging and the neural processes that regulate it to effectively treat age-related visual dysfunction and disease linked to neural dysfunction.

*Results:* Mitochondria are cellular organelles that enable energy metabolism, and essential cellular signaling processes. Mitochondrial function declines with age in the visual system and is linked with the development of age-related visual disease. Here, the authors present evidence that improving mitochondrial function can treat age-related visual decline. They report that a loss of acuity emerged in mice near 18 months (early old age) and declined with age until 34 months (extreme old age) when it was reduced by ∼60%. Daily administration of the mitochondria-acting peptide, SS-31, from 18 months largely prevented subsequent age-related visual decline. Application from 24 months, when moderate visual impairment was present, led to complete recovery of visual acuity within 2 months, which persisted for at least 3 months after SS-31 was withdrawn. A single dose at 24 months was also able to delay visual decline. Moreover, 2 months of SS-31 administration from 32 months, after much more severe visual dysfunction was manifest, substantially improved function by 34 months.

*Implications and future directions:* The study reveals that spatial measures of visuomotor function can identify age-related visual decline in mice that is largely preventable and reversible early in its course, by treatment with a mitochondrial-acting peptide. That visual dysfunction late in life is partially reversible with the peptide, also indicates that treating mitochondrial dysfunction has the potential to provide a benefit at any age. In addition, that restored function endured after the peptide was withdrawn indicates that improving mitochondrial function elicits long-lasting beneficial changes in the aging visual system. By linking mitochondrial dysfunction with visual aging, the data also suggests that improving mitochondrial function is a promising approach for treating age-related visual disease.

## INTRODUCTION

Visual decline related to normal aging contributes to disability and reduces health span (Owsley 2011; 2016). Hardening (presbyopia; Lafosse et al, 2020) and clouding (cataracts; Asbell et al, 2005) of the lens are common optical consequences of aging that can seriously impair visual function. The development of devices and procedures (eg. corrective eyewear; lens replacement surgery, etc.) to reduce refractive error in these conditions have drastically reduced the burden of age-related optical visual impairment in the world, though it is still substantial (Holden et al, 2014). Independent of optical factors, untreatable decline of vision due to age-related neurological dysfunction is also a major source of impairment (Owsley, 2011). This can present as the deterioration of spatial visual function (Elliot, 1987), luminance modulation (Woi et al, 2016), binocular processing (Arani et al, 2019), color perception (Karwatsky et al, 2004), sensitivity to motion (Habak and Faubert, 2000), and dark adaptation (Gaffney et al 2012; Jackson et al, 1999), among other impairments. Some visual functions, such as blur adaptation, however, appear to remain intact with age (Elliot et al, 2007). Age also predisposes the visual system to develop age-related diseases, such as glaucoma, diabetic retinopathy (DR), and age-related macular degeneration (AMD). These blinding diseases are common-approximately one in three elderly persons has some form of vision-reducing eye disease by the age of 65 (NEI, Eye Health Data and Statistics)- and have risen with medical advances that have extended lifespan. Unfortunately, neither the etiology of age-related visual decline, nor the mechanisms of how age contributes to blinding diseases, are sufficiently well understood to enable effective treatment. Thus, despite the prospect of increased longevity, the elderly face reduced quality of life with increased risk of disability from falls, immobility and depression linked to visual impairment (Court et al, 2014).

Since age itself contributes to untreatable visual decline, understanding the natural history and pathophysiology of visual aging has clinical relevance. The typically large individual-to-individual variability in the effects of aging on human vision, and the small sample sizes that are often used in clinical studies, however, are not optimal conditions for identifying fundamental pathophysiology of visual aging. Alternatively, using inbred mice in visual aging studies, with their reduced subject-to-subject variability, has the potential to characterize the natural history of mammalian visual aging, and help identify its neurophysiological substrates. Whereas there are reports of declining visual acuity with age in mice (Weber et al, 2015), to our knowledge, a careful quantification of spatial visual function-acuity and contrast sensitivity-among the most widely used and clinically-relevant measures of visual function-across the lifespan has not been completed.

Characterizing visual decline over the lifespan may also aid in understanding and modeling age-related retinal disease. Despite the link between age and disease, preclinical models of sporadic age-related blinding diseases have not routinely included advanced age as a variable. Rather, they have focused on genetic mutations or physiological modifications that induce retinal degeneration in younger animals. Whereas this approach has deepened our mechanistic understanding of retinal degeneration, it has not necessarily advanced relevant rodent models of age-related blinding diseases-particularly for sporadic AMD-, which is often modeled with mutations linked to inherited forms of disease like retinitis pigmentosa (e.g. The Royal College of Surgeons rat). The lack of age as a variable in preclinical models may have contributed to the problem that few interventions have been successfully translated for use in human age-related visual disease. Thus, understanding the contributions that age makes to visual decline, is likely an important step in determining what distinguishes age-related visual decline from age-related visual disease.

Numerous abnormalities in cellular physiology have been linked with aging (Takahashi et al, 2000; López-Otin et al, 2013), but among the most evidential is that compromised mitochondrial bioenergetics is a regulator of age (Kauppila et al, 2017) and disease (Annesley and Fisher, 2019). Such dysfunction is measured as reduced ATP production, decreased membrane potential, oxidative stress, organelle swelling, cristae damage, and decreased DNA copy, among other deficiencies. Since mitochondria are present in all cells, and mitochondrial function is required for a host of cellular functions-regulation of apoptosis, calcium buffering, nuclear genome signaling, reactive oxygen species (ROS) production, steroid synthesis, immune system signaling, cell cycle and cell growth regulation, etc. (Bock and Tait, 2020; Tait and Green, 2012; Fang et al, 2016), it is not surprising that the consequences of mitochondrial dysfunction would be diverse and affect a wide variety of physiological processes and organ functions. Consistent with this are reports linking dysfunctional mitochondria to aging (Bratic and Larsson, 2013) and a wide variety of diseases, including those that are age-related, such as cancer, metabolic disease and diabetes, inflammatory disease, neuropathy, nephropathy, and in neurodegenerative conditions like Alzheimer’s, Parkinson’s, and Huntington’s disease (Swerdlow et al, 2017; Lin and Beal, 2006). In the visual system, mitochondrial mutation-linked dysfunction subserves Leber Hereditary Optic Neuropathy, which is the most common inherited mitochondrial disorder (Pilz et al, 2017). In addition, mitochondrial dysfunction, in the form of impaired mitochondrial dynamics (fusion and fission) has also been linked to primary open angle glaucoma (Shim et al, 2016), and dysfunction has been linked to AMD (Nashine et al 2019; Ferrington et al, 2016).

One way to test the hypothesis that mitochondrial dysfunction is a pathophysiology of age-related visual decline would be to treat aged mice with a pharmacological agent that targets mitochondria and improves mitochondrial function, and evaluate its ability to prevent or reverse visual decline. If the treatment were beneficial, it would provide evidence supporting mitochondrial dysfunction as a pathophysiology of age-related visual decline, and by association, implicate mitochondrial dysfunction in age-related visual disease. In addition, a therapeutic target and a potential therapeutic agent to treat human age-related visual decline and disease, would have been identified. SS-31 (A.K.A Elamipretide, MTP-131; Zhao et al, 2003; Schiller et al, 2000) is among the most promising candidate compounds to test a mitochondrial hypothesis of visual aging. It is a synthetic tetrapeptide with an alternating aromatic-cationic motif that is attracted to the negative surface charge of mitochondrial membranes enriched in cardiolipin (Berezowska et al, 2003), where it alters biophysical properties through electrostatic and hydrophobic interactions (Birk et al, 2013). Other work has linked SS-31 treatment effects to the function of the electron transport chain (Chavez et al, 2020) and alterations of the biophysical properties of cardiolipin-enriched membranes (Mitchell et al, 2020). Cardiolipin plays a central role in the formation of crista, the electron transport chain, and in regulating the function of ETC complexes and ATP synthase (Mileykovskaya and Dowhan, 2014). In addition to its antioxidant activity (Zhao et al, 2005), SS-31 also mitigates peroxidation of cardiolipin and inhibits cytochrome *c* peroxidase activity (Birk et al, 2014; Birk et al, 2013). The combined antioxidant and apparent mitochondrial rejuvenating properties of SS-31 appear to benefit age-related decline in skeletal muscle (Siegel et al, 2013; Campbell et al, 2018), kidneys (Sweetwyne et al, 2017), and brain (Tarantini et al, 2017). In age-related disease models, SS-31 has been effective at ameliorating dysfunction associated with hypertension (Dai et al, 2011), heart failure (Dai e al, 2013), and glaucoma (Wu et al, 2019). SS-31 treatment is also effective at treating visual decline in a mouse model of diabetic retinopathy (Alam et al, 2015a), which has linked mitochondrial dysfunction to visual disease. Thus, in the study presented here, we tested the hypothesis that SS-31 would prevent and reverse spatial visual decline in a mouse model of visual aging.

## MATERIALS AND METHODS

### Animal subjects

Experiments complied with the policies of Weill Cornell Medicine Animal Care and Use Committee. 224 C57BL/6 mice of both sexes ranging from ∼1-34 months of age were obtained from National Institute of Aging aged rodent colonies curated by Charles River Laboratories, and from Charles River Laboratories directly, and were group housed at the Burke Neurological Institute vivarium. They had *ad libitum* access to food (Rodent Diet 5053) and acidified water, were maintained at 68°-76°F with 30-70% relative humidity, and with a photoperiod of 12 h light (06:00 lights on) / 12 h dark (18:00 lights off).

### SS-31 administration

Mice in most experiments were injected subcutaneously (s.c.; once/day with a 1 mg/kg solution of the tetra-peptide SS-31, provided by Stealth Biotherapeutics, Newton, MA, USA) dissolved in 0.9% sterile saline (pH 5.5-6.5), or were injected with 0.9% saline alone. In some experiments, mice were administered SS-31 once/day as an ophthalmic-formulated solution (Ocuvia; provided by Stealth Biotherapeutics, Newton, MA, USA) daily via eye drops (in 0.01 M sodium acetate buffer solution (pH 6.00); 5 μl/eye), or buffer alone.

### Tests of spatial and temporal visual function

Spatial frequency and contrast thresholds for opto-kinetic tracking of sine-wave gratings were measured using a virtual opto-kinetic system (OptoMotry, CerebralMechanics Inc, Medicine Hat, Alberta, Canada; Prusky et al., 2004; Douglas et al, 2005). Vertical sine-wave gratings projected as a virtual cylinder and drifting at 12°/s, or gray of the same mean luminance, were displayed on four computer monitors arranged in a square around a small elevated platform. For testing, a mouse was placed on the platform and allowed to move freely. The hub of the cylinder was then centered between the animal’s eyes as it shifted its position in order to fix the spatial frequency of the grating. Gray was projected when the mouse was ambulating, and a grating was projected when it was stationary, which when visible to the animal, elicited tracking movement of the head and neck. The presence of tracking under each stimulus condition was appraised via live video with a yes/no criterion by an observer blind to the group identity of the mice, and a threshold for tracking was established using a method of limits procedure. A spatial frequency threshold (acuity)- the highest spatial frequency to elicit tracking of a grating at maximal contrast) through each eye (Douglas et al, 2005)- was obtained in a testing session in a few minutes. In some sessions, spatial frequency and contrast thresholds (lowest contrast to elicit tracking) at six spatial frequencies (0.031, 0.064, 0.092, 0.103, 0.192, 0.272 c/d to generate a contrast sensitivity function (CSF)) through each eye separately (by changing the direction of stimulus rotation) were measured (14 thresholds), in ∼30 min. Michelson contrast sensitivity was calculated from the contrast thresholds using the average screen luminance (maximum−minimum)/(maximum+minimum). Experimental animals and their controls were assessed in the same testing session. Thresholds for all animals were obtained under photopic lighting conditions (screen luminance=54 lux), which selectively measures cone-based visual function (Alam et al, 2015b). Photopic measures commenced at one month of age; an age at which mature function is normally established (Prusky et al, 2004). Since it was not feasible to measure the same cohort of mice over their entire life, multiple cohorts that overlapped in age were employed, which enabled the sampling of visual function over the span of adult mouse life. Rod-based function under scotopic conditions (screen luminance =1 lux; Alam et al., 2015b) was assessed in some animals after they were individually dark adapted (>6 h). For this, ND filters (Lee Filters) were placed over the monitor screens (6.9ND), the testing arena was made light tight, and a near infrared-sensitive camera with near infrared lighting (Sony Handycam DCR-HC28, Sony, Japan) was used to image the animals.

In order to determine whether spatial visual decline with age, and its remediation with SS-31, were accompanied by changes in temporal neural processing, we also investigated whether photopic temporal frequency tracking thresholds-a behavioral analog of neural temporal processing (Umino et al, 2008)- was affected by age and SS-31 treatment. For these experiments, the stimulus temporal frequency was maintained at 1.5 Hz; a temporal frequency at which it was feasible to assess tracking responses across a wide range of stimulus speeds. A spatial frequency threshold was then measured in vehicle-treated mice at 3 and 26 months, and in mice at 26 months treated daily from 24 months of age with either s.c. injections of SS-31 or vehicle.

### Ophthalmic assessments

Optical clarity was evaluated with a visual inspection. A biomicroscope (slit lamp) or dissecting microscope was used in some cases to inspect the cornea for clarity, size, surface texture and vascularization, and the iris was evaluated for pupil size, constriction, reflected luminescence and synechia. On some occasions, pupils were dilated with a drop of 0.05% tropicamide ophthalmic solution, and the lens was inspected for cataract with an indirect ophthalmoscope or a dissecting microscope (Merriam and Focht, 1962; Worgul et al, 1993). At the same time, the fundus was inspected for damage, degeneration, retinal vessel constriction and optic nerve head abnormalities. The data from animals with reduced optical clarity or fundus abnormalities were excluded from the analysis.

### Statistical analyses

Two-way, repeated-measures ANOVAs were used to make group comparisons using the statistical software package Prism. Post-hoc multiple comparisons were performed using the Tukey’s or Bonferroni correction methods. Statistical comparisons were considered significantly different at *P*<0.05. Simple linear or segmental nonlinear regression fit analyses were used to compare slopes.

## RESULTS

### Spatial visual function declines with age

Fig. 1A shows that acuity averaged ∼0.39 c/d from 1 to 18 months of age. This value is consistent with previous reports of normal adult visual function using the same strain, apparatus and measurement procedures (Prusky et al, 2004; Douglas et al, 2005; Prusky et al, 2006; Tschetter et al, 2013, Alam et al, 2015b). Function gradually declined after 18 months to ∼70% normal adult function at 34 months (0.39 c/d Vs 0.12 c/d; p<0.01); the oldest cohort tested.

**Figure 1:**
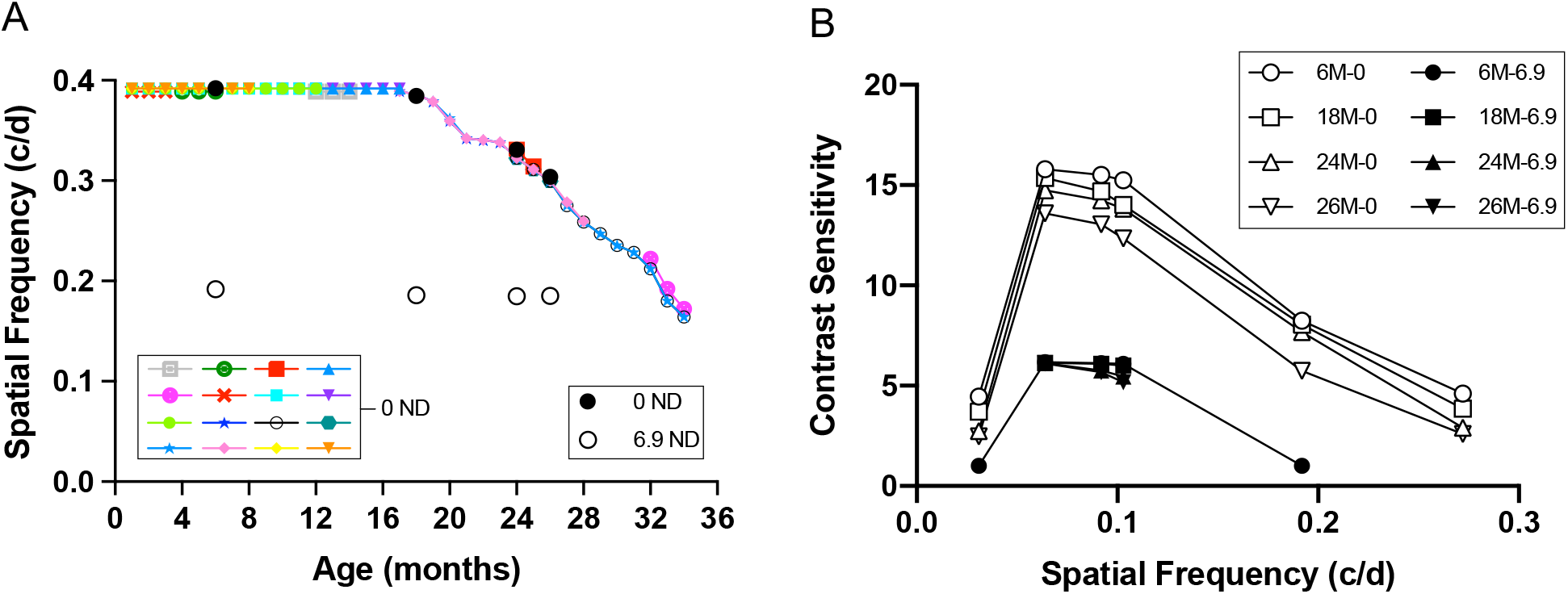
Mouse spatial visual function declines with age. A; Spatial frequency thresholds. Colored symbols depict weekly average photopic measures of separate cohorts of mice (0 ND panel; 16 cohorts; total n=224). Function near 0.39 c/d was maintained from 1-18 months, and declined thereafter with periods of increased and decreased rates of change. Function at 34 months (0.16 c/d) was reduced by 59% relative to normal adult values. Vertical standard deviation lines are depicted in this and other panels, but they are often smaller than the data symbols. A cohort of mice (n=6) was measured under both photopic (black closed circles; 0 ND) and scotopic (white open circles; 6.9 ND) conditions. Photopic acuity showed substantial decline with age (6-26 months= −22.4%) in the group, scotopic acuity varied little over the same period (−2.8%). B; Contrast sensitivity functions. Groups of mice (6-26 months of age; symbols identified in the legend) were measured under photopic (0 ND) and scotopic (6.9 ND) conditions. Photopic function generally declined with age at all spatial frequencies. Scotopic values showed little evidence of change with age at spatial frequencies near peak sensitivity.

One group of mice was measured periodically between 6 and 26 months under both photopic and scotopic conditions, the results of which are also presented in Fig. 1A. Scotopic thresholds at 6 months averaged 0.19 c/d - characteristically lower than photopic thresholds (Prusky et al, 2004; Douglas et al, 2005; Alam et al, 2015b) - and varied little out to 26 months (scotopic 0.192c/d to 0.187c/d, n=6, p<0.0001); photopic 0.39c/d to 0.30c/d, n=6, p<0.0001). This reveals that normal scotopic function was maintained over the same period that photopic function was in decline. The trend of declining photopic function and more stable scotopic function continued in measurements of contrast sensitivity (CS) of mice aged 6-26 months, which is illustrated in Fig. 1B. The contrast sensitivity functions (CSFs) of 6 month old mice were consistent with previous work on the same strain and with the same measurement procedures (Prusky et al, 2004; Douglas et al, 2005; Alam et al, 2015^b^). Photopic function declined gradually across spatial frequencies from 6-24 months, and decline accelerated from 24-26 months. Scotopic CS decreased little as a function of age. Only spatial frequencies near peak sensitivity were measurable by 18 months, and thus, a characteristic convex function could not be obtained. At maximum sensitivity, a modest decline was evident at 24 months, relative to 6 months old.

### SS-31 treatment can slow or reverse age-related visual decline

After characterizing a decline in photopic acuity and CS with age, we set out to determine whether SS-31 would prevent the advancement of age-related visual impairment. For this, mice were treated daily from 18 months with s.c. 0.9% saline, or SS-31 (1mg/kg). Photopic spatial frequency and contrast thresholds were measured regularly in the same animals until 24 months. Fig. 2A shows that the SS-31-treated cohort’s photopic acuity was substantially preserved; the rate of decline was slower in SS-31 treated-than in placebo-treated mice (SS-31= −0.0008097; Placebo= −0.002485). Indeed, normal function was maintained for at least 10 weeks after treatment, and thereafter, the rate of decline was slower than in placebo controls. Compared to baseline measures at 18 months (0.39c/d), by 24 months (0.32c/d), the SS-31 cohort declined by 6% compared with 16% in the placebo cohort. Fig. 2B shows that a similar trend of SS-31 preserving function with treatment from 18 months was evident in CS measurements, but the effect was not as straightforward to interpret; in general SS-31 treatment led to a modest improvement of function, but at some spatial frequencies, treated animals were not different than controls. Together with the results of acuity measures, these data indicate that the treatment benefit of SS-31 on spatial vision was at the high spatial frequency range.

**Figure 2:**
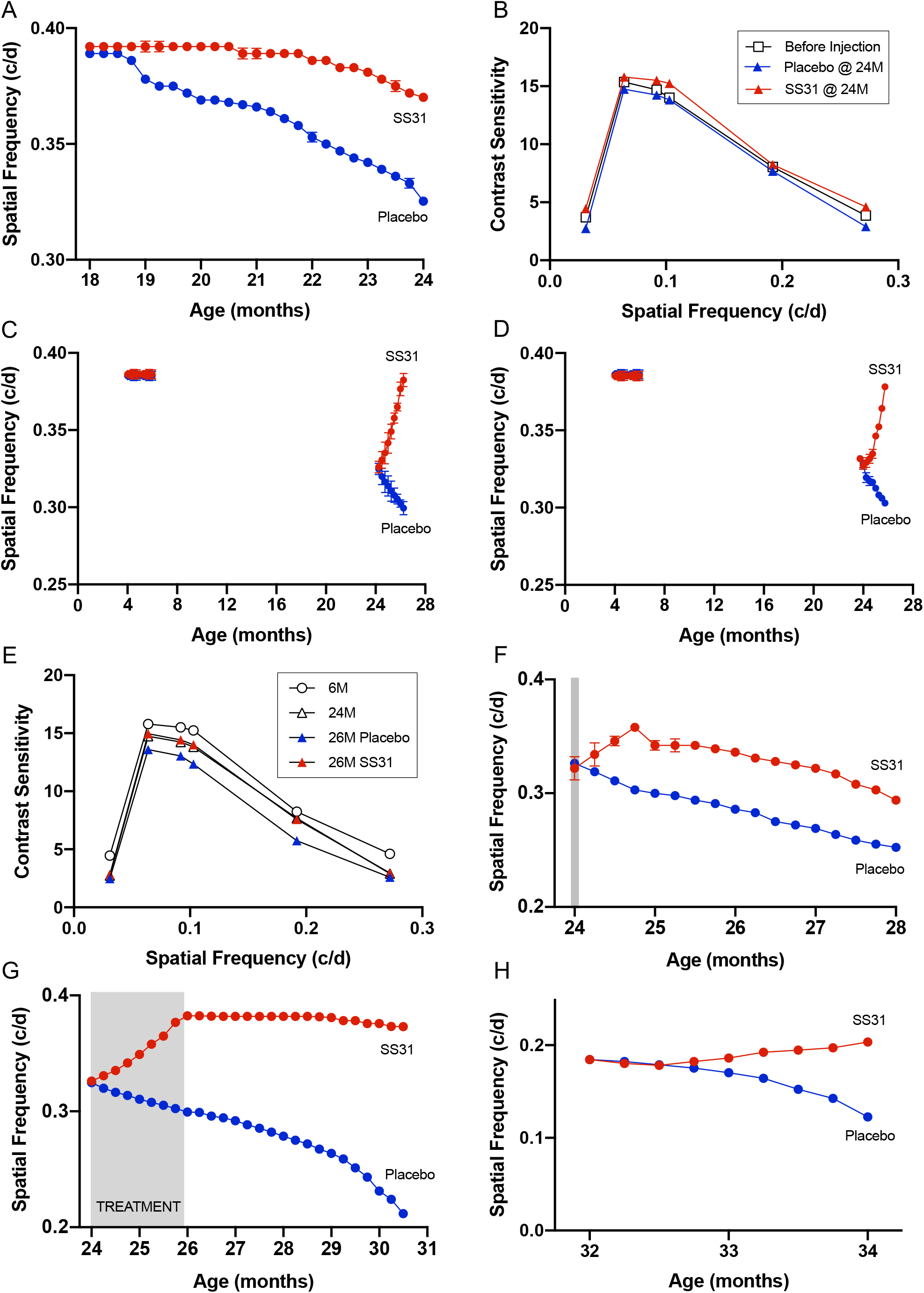
SS-31 can treat age-related photopic vision loss using various dosing durations and methods of administration – even in mice of extreme old age. A; Spatial frequency threshold. Treatment from 18 months (red circles; n=6) enabled the maintenance of function within a normal range for more than 2 months, and thereafter, reduced the rate of decline relative to placebo-treated group (blue circles; n=6; 5.5% Vs 16.3% decline at 24 months). B; Contrast sensitivity. Treatment from 18 months (red triangles) led to improved function relative to placebo-treated animals (blue triangles), and resulted in better function at most spatial frequencies than that of untreated animals before treatment was initiated (open squares). C; Effect of daily SS-31 s.c. treatment on spatial frequency thresholds. Drug-treated mice (n=10; red dots) from 4-6 months were similar to placebo treatment (n=10; left side; 0.39 Vs 0.39 c/d; blue dots). The same treatment from 24 months (n=52) reduced decline within 1 week, and restored normal function by 26 months (0.38 c/d), whereas placebo-treated animals (n=49) continued to decline (0.30 c/d at 26 months). D; Effect of daily SS-31 eye drop treatment on spatial frequency thresholds. Drug-treated mice (red dots) from 4-6 months (n=10) were similar to placebo treatment (n=10; left side; 0.39 Vs 0.39 c/d; blue dots). The same treatment from 24 months (n=52) reversed decline after 4 weeks and restored normal function by 26 months (0.38 c/d), whereas placebo-treated animals (n=49) continued to decline (0.30 c/d at 26 months). E; Effect of daily eye drop SS-31 treatment on contrast sensitivity. Drug treatment (red triangles) maintained visual function at 24 month norms (open triangles), relative to placebo treatment (blue triangles), but did not restore function to 6 month norms (open circles). F; Effect of a single SS-31 s.c. treatment at 24 months on photopic spatial frequency threshold (acuity; red dots; n=7)). Function improved for 4 weeks after treatment and then paralleled age-related decline in placebo-treated mice (blue dots; n=7) out to 28 months of age (0.28 Vs 0.25 c/d). G; Effect of withdrawing treatment (daily s.c.; gray shading) from 24-26 months of age. Near-adult normal function was maintained in SS-31—treated animals (red dots; n=8) for more than 4 months after withdrawal at 26 months, and much slower loss of function with age was exhibited thereafter relative to placebo controls (blue dots; n=8; 0.37 Vs 0.21 c/d). H; Treatment benefit of SS-31 on spatial visual impairment in advanced age. Daily SS-31 eye drop treatment (gray shading) from 32 months of age led to improved spatial frequency thresholds within 2 months (0.18 Vs 0.20 c/d) compared with loss of function in placebo-treated mice (0.18 Vs 0.12 c/d). Standard deviation is plotted as vertical bars on all panels but is occluded by symbols in some.

We then investigated whether photopic visual function could be restored in older, 24 month-old mice, an age at which a degradation of 18% of the photopic acuity, and 5% of photopic CS at peak sensitivity, was present (see Fig. 1). 24 month-old mice were treated with daily s.c. placebo injections, or SS-31, for two months. Fig. 2C shows that acuity was improved relative to placebo within 1 week of initiating treatment (p<0.0001), and continued to improve thereafter. After 8 weeks of treatment, the threshold of the SS-31 cohort returned to pre-decline values (p>0.9999). Young mice that underwent the same course of treatment did not change (p>0.9999), showing that the application of SS-31 did not augment normal function.

In an effort to target the eye more directly and to explore the feasibility of using eye drops to deliver a therapeutic dose of SS-31, we replicated the two-month course of vehicle or SS-31 treatment in 24 month-old mice, with eye drops. Fig. 2D shows that whereas the eye drop application of SS-31 proved slower to improve function than s.c. application (5 Vs 1 week), the treatment was able to restore the photopic acuity to the same degree as s.c. delivery over the course of 2 months. Fig. 2E shows that the CSF of aged mice with eye drop delivery of SS-31 for 2 months was maintained at 24-month pretreatment baseline values.

In order to investigate the potency and persistence of SS-31 to reverse age-related visual decline, different treatment durations were employed and acuity was measured in the same animals for the subsequent 2 - 4.5 months. Fig. 2F shows data from 24 month-old mice that were treated with a single subcutaneous injection of placebo, or SS-31, and followed with repeated measures for 4 months. The treatment led to improved function within 1 week (p<0.0001), which continued to improve for 3 weeks. Thereafter, function declined at a rate that closely resembled loss of function in the vehicle-treated group up to 28 months (SS-31= −0.00395; placebo= −0.005218). In another experiment, mice were treated from 24-26 months with daily s.c. injections of SS-31 or placebo from 24 months (treatment that led to full recovery of function), after which treatment was suspended. Figure 2G shows that as late as 4.5 months after treatment was suspended (∼30 month-old mice), the SS-31 cohort maintained 97% of their recovered function, whereas the placebo group had declined to 46% of normal adult function.

Whether SS-31 was able to improve visual function in extreme old age, when much more visual impairment was present, was also investigated. At 32 months, when ∼50% loss of visual function had occurred, mice were treated with daily subcutaneous injections of placebo or SS-31 for 2 months. Figure 2H shows that although there was little deviation in age-related visual decline with SS-31 treatment for the first 2 weeks, function was gradually improved thereafter. By the last measurement at 34 months, the SS-31 cohort had improved significantly (p< 0.0001), whereas the placebo group had declined to 31% normal function (p< 0.001; SS-31 vs placebo at 34 months p< 0.001). This indicates that once recovery commenced in the 32-month old SS-31 treated group, the rate of improvement was slower than in animals with the same 2-month treatment regimen begun at 24 months.

Several SS-31 administration schedules were used in the study to establish that the peptide was efficacious at reversing age-related visual decline, and to gain insight into the kinetics of the drug’s action. In order to enable direct comparisons between various treatment regimens, Fig. 3A plots results of experiments shown in Fig. 2C, F, G and H on the same time scale to enable direct comparisons of the rate and magnitude of SS-31 responses on acuity with treatment for 2 months or more. This revealed that whereas daily treatment from 24 months resulted in the greatest benefit, the initial rate of improvement in function was faster when a single treatment, or weekly treatments, were used. The rate of recovery with daily treatment from 24-26 months (slope = 0.007310) was also faster than the recovery with treatment from 32 months (slope = 0.002830).

**Figure 3:**
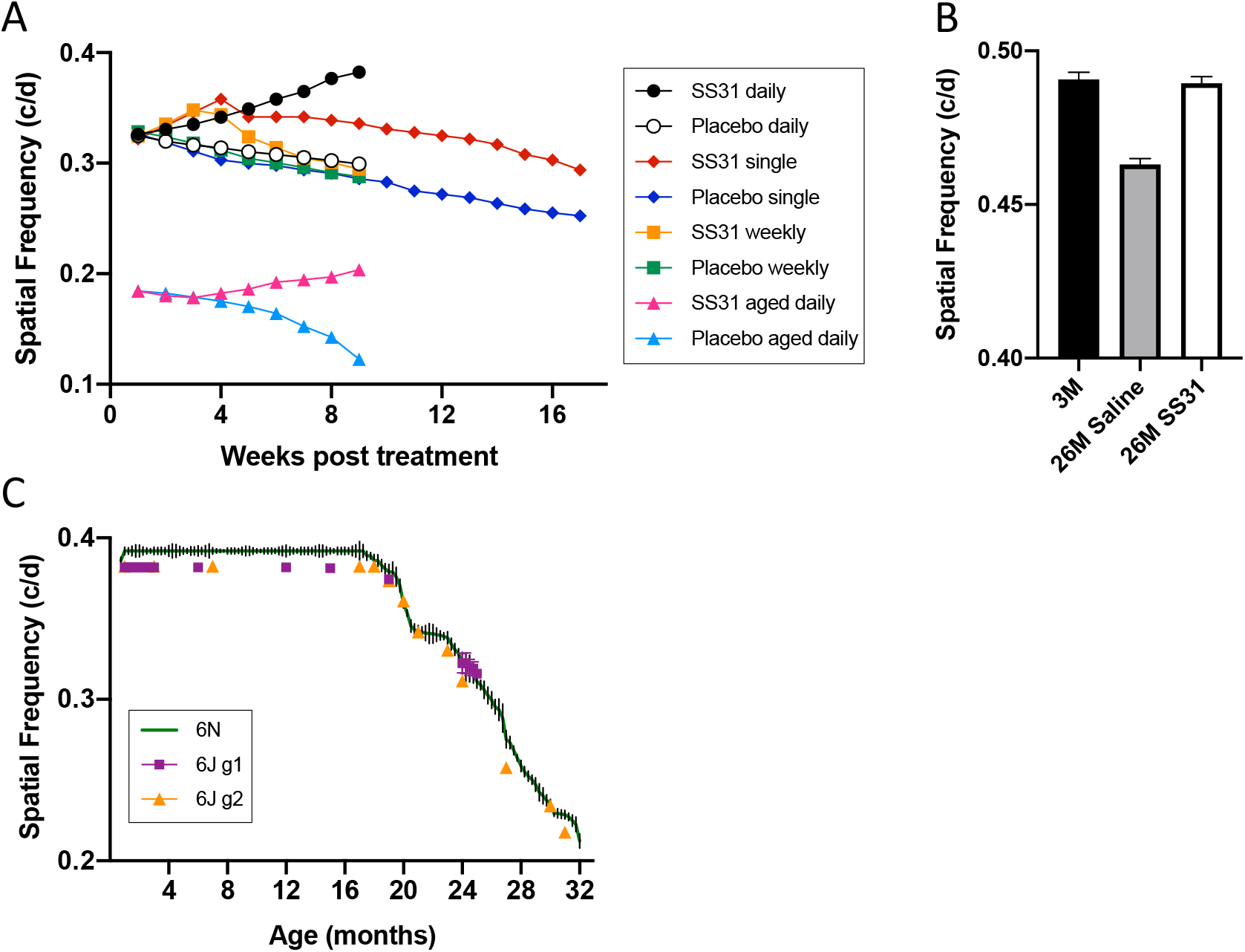
Additional analyses and experiments. A; Comparison of SS-31 treatment regimens; spatial frequency threshold results shown in Figs. 2C, F, G and H plotted from start of treatment (groups labeled in legend). Treatment from 24 months (black filled circle=SS-31; white circle=placebo control) produced the largest benefit, the initial rate of improvement with treatment at 24 months appeared better when a single treatment (red diamond=SS-31; blue diamond= placebo control), or weekly treatments (orange square=SS-31; green square=placebo control), was used. The rate of recovery with daily treatment from 24-26 months (black circle) was faster than the recovery with treatment from 32 months (pink triangles=SS-31; blue triangles=placebo control). B; Photopic temporal frequency thresholds of untreated mice at 3 months (black bar), 26 month old mice treated with saline (control) from 24-26 months (gray bar), and 26 month old mice treated with SS-31 from 24-26 months (white bar; vertical standard deviation lines plotted on each bar). The results show temporal processing declines with age, and can be restored to normal adult values with SS-31 treatment. C; Comparison of age-related decline in C57Bl/6N (plotted as vertical standard deviation bars from Fig. 1A) and C57Bl/6J mice plotted as colored symbols for each cohort tested (standard deviation lines are often occluded by symbols). Adult function (1-18 months) of C57Bl/6J mice was slightly lower than in C57Bl/6N mice, but the pattern of age-related visual decline was similar.

We investigated whether temporal frequency thresholds were altered by age and SS-31 treatment. Fig. 3B shows that the threshold responses of vehicle-treated 26-month old mice were lower than in 3-month old mice (p<0.001), and that SS-31 treatment from 24-26 months was able to normalize the function (3 month vs 26 month + SS-31, p= 0.6114). This shows that temporal processing of photopic information, in addition to spatial processing, declines with age, and can be improved with SS-31 treatment.

The core results in this study were generated using C57Bl/6N mice acquired from NIA colonies. C57Bl/6N mice have been reported to carry an RD8 mutation of the Crb1 gene that can lead to a retinal degeneration phenotype (Mattapallil et al, 2012). The C57Bl/6J strain, on the other hand, has been reported to carry a 5 exon NNT deletion, but does not appear to normally carry mutations, such as RD8 (Ronchi et al, 2013). In an effort to determine whether age-related visual decline in C57Bl/6 mice varied by sub-strain and/or retinal mutation, we compared the visual function of 2 cohorts of C57Bl/6J mice raised in-house as they aged, acquired from The Jackson Lab, with the results generated in C57Bl/6N abstracted from Fig. 1A. Fig. 3C shows that whereas the baseline adult function of C57Bl/6J mice was lower than that in the C57Bl/6N strain, the overall pattern of age-related visual decline was comparable. This indicates that the age-related decline of visual function we report here is likely a characteristic of the C57Bl/6 strain, and not the result of a known mutation that affects the function of the retina.

## DISCUSSION

We investigated how age affects spatial vision in C57Bl/6 mice; the most widely used mouse strain in neurobiology research. We report that acuity measured in photopic luminance conditions was normal up to ∼18 months of age, and declined gradually thereafter, until advanced old age (34 months) when <40% normal function remained. Little change in acuity measured under scotopic luminance conditions occurred over the same time span. Previous studies have reported that SS-31 targets mitochondria, due to its affinity for cardiolipin-rich membranes, and has a beneficial effect on mitochondrial function. These properties enabled us to use SS-31 as a tool to investigate whether mitochondrial dysfunction contributes to age-related visual decline, and whether improving mitochondrial function can treat it. We generated evidence that SS-31 was effective at treating (both preventing and restoring) photopic, age-related loss of spatial vision, and at improving the ability of the visual system to process photopic temporal information. The results of the study provide a framework to investigate the cellular and molecular substrates of how mitochondrial dysfunction with age leads to visual decline, and how improving mitochondrial function enables improvement of visual function. They also provide an impetus to investigate whether mitochondrial dysfunction is a treatable pathophysiology of human visual aging and age-related visual disease.

### Decline of spatial vision with age reflects cone-mediated function

Measures of spatial vision-acuity and contrast sensitivity-report how effectively neurons in the visual system process size and luminance, respectively, in a visual scene. Since changes in spatial visual thresholds reflect changes in the structure and function of the visual system, one of the goals of the study was to identify neural circuitry that underlies age-related visual decline. To this end we identified a consequential role for cone circuitry in spatial visual decline with age. We report that over more than half of the normal murine lifespan, -up to ∼18 months of age-photopic and scotopic spatial visual function was stable; in the face of advancing age, the mouse spatial visual system is able to maintain normal function. The physiology of this resiliency was not investigated in the study. Near 18 months of age, photopic visual function commenced a gradual decline, with ∼18% deterioration of acuity by 24 months. Proportionally little change was measured in scotopic function up to 26 months.

These results do not imply that photopic changes in spatial vision are the only behavioral changes in the mouse visual system function with age, or that rod-based function is spared the effects of aging. Indeed, there are reports of age-related changes in rod structure and function in mice (Kolesnikov et al, 2010; Rohrer et al, 2003; Wang et al, 2018). There are a number of possible explanations as to why scotopic spatial vision was not substantially affected by age in the study. One possibility is that the scotopic spatial visual thresholds are normally much lower than photopic thresholds, and as such, it might be more difficult to measure significant changes; normal adult acuity measured under scotopic conditions (∼0.2 c/d) is ∼50% lower than that measured under photopic conditions changes (∼0.39 c/d) Thus, there may be proportionally less headroom to measure scotopic changes in spatial vision with age. We think this explanation is unlikely. Indeed, we did measure reductions in scotopic thresholds with age, but they were quantitatively much smaller than the changes in photopic function. This is despite the fact that variability in the measurement of scotopic and photopic measures was similar, and therefore, there should have been a similar ability to measure change if it was present.

A more likely explanation is that photopic and scotopic vision serves different purposes for mammals, and that spatial vision is more dependent on cone circuitry. The measures we made are not likely to be a sensitive behavioral test of rod functional decline with age. In humans, rod-driven function is known to decline with age before cone-driven function, which is often manifest as a deterioration in peripheral (rod-biased), not foveal (cone-biased) function, and in impaired dark adaptation (Jackson et al, 1999). Thus, the reduced effect of age on scotopic function in our study probably reflects the low-resolution nature of the rod spatial visual system due to the neural pooling of rods, which serves a different function than cone-based circuitry. Certainly mice, even though they are nocturnal and do not possess a fovea, do not rely on rod-based vision for competent spatial visual function, and instead use a higher resolution cone-based system. Thus, our spatial measures, even though they were made using opto-kinetic responses that do not depend on cortical visual circuits (Douglas et al, 2005), were likely biased toward the detection of functions associated with cone-based circuits, including those that are downstream of the retina. It is possible that different behavioral measures that rely on a different-possibly cortical-functions, such as the Visual Water Task (Prusky et al, 2000) would reveal greater rod-based decline of spatial function with age.

### Restoration of Cone-Mediated Spatial Function with SS-31 Treatment

The study reveals behavioral evidence that SS-31 treatment rather selectively improves age-related loss of photopic function, which links the effect to mitochondrial function; at least to the extent that treating animals with the peptide that has a selective affinity for mitochondria, and that SS-31 has been shown to improve mitochondrial function. These results make the study of clinical relevance, because age-related visual decline becomes most debilitating in humans when cone function linked to spatial vision becomes involved. Thus, the study may provide a working model of this phenomenon. Indeed, our study, lends support to the prospect that the most debilitating effects of age-related visual decline and disease, may be treatable. SS-31 is in clinical trials to evaluate its ability to ameliorate vision loss in humans (ReCLAIM-2-https://www.stealthbt.com/wp-content/uploads/1907_StealthBio_Phase2_Infographic_FINAL-converted-1.pdf; ReSight-https://www.stealthbt.com/wp-content/uploads/StealthBio_ReSIGHT_4.24.2018.pdf), and further studies can now be justified to investigate this possibility. In addition to the ability of SS-31 to treat age-related visual decline in this study, it is noteworthy that we found no evidence in young (control) mice that SS-31 treatment leads to super-normal function; many therapeutics do not display this feature, which is a common safety concern. Thus, the ability to treat dysfunction without compromising normal function, also lends preclinical support to the safety profile of SS-31.

The evidence that SS-31 can slow and reverse age-induced decline in visual acuity and contrast sensitivity of aged mice is supported by multiple lines of evidence. This includes the finding that improvements in SS-31-treated cohorts (relative to placebo) occur over a range of administration start times, treatment durations, and types of drug delivery. It also includes the intriguing finding that SS-31 treatment is able to improve function in mice even at a very advanced age (32 months), albeit at a slower rate than in younger aged animals. We did not investigate the effect of age and SS-31 treatment on temporal neural processing in the visual system as extensively as we did spatial processing. However, the finding that temporal neural processing declines with age up to 26 months, and can be normalized with SS-31 treatment, indicates that the benefits to the visual system of improving mitochondrial function are pervasive. Not only do these convergent results bolster confidence in our conclusion that SS-31 can prevent and reverse age-related spatial visual decline, they also indicate that SS-31 could be compatible with a wide variety of treatment strategies and ages in humans.

The persistence of the SS-31 treatment benefit after withdrawal-even following a single treatment- is a particularly provocative finding of the study. It indicates that in addition to SS-31 affecting the function of the visual system while it is biologically active-on the order of hours-it appears able to alter the physiology of the visual system in an enduring and beneficial way. This result and the finding that the schedule of treatment can affect the rate of recovery and the rate of decline of the treatment effect provides both a challenge and an opportunity for translational studies. A challenge because the effects do not appear to follow traditional expectations of pharmacokinetics; an opportunity because SS-31 treatment may be compatible with a wide variety of treatment strategies. Indeed, the data indicate that it may be possible to commence an SS-31 treatment regimen with a relatively high ‘loading dose’ to propel improvement of function, and then maintain the improvement with lower dose or a more dispersed schedule of treatment. More preclinical studies would likely be necessary to develop a treatment regimen that maximizes the benefit of SS-31 treatment, while reducing the drug exposure to a minimum.

### Implication of the Study Results for Future Mechanistic Studies

The present study was not designed to identify the cellular mechanisms of aging, nor to determine how, or where, mitochondrial function and action may be involved in the visual aging process and its remediation. Instead one of the aims of the study was to generate a detailed description of how age affects spatial visual function. To that end, we present evidence derived from >200 mice across multiple cohorts, that visual aging can be quantified in a way that models the human visual aging process. This is a significant advance because there is now a framework for future studies to test hypotheses regarding the cellular mechanisms of visual aging, and to interpret the results in terms of the rate of change over time; not only at single age points. For example, if an experiment compared the spatial vision of young mice (e.g. 3 months old) to mice of advanced age at 12 months or even 18 months of age, our data indicate that one would not expect to observe any differences, since spatial vision appears to be intact until 18 months. In addition, our data reveal that comparisons between control groups at 18 months of age, and older animals, may be a more informative experiment than comparisons with young adult mice (e.g. 3 months), since it would reduce the influence of spurious changes that occur with age between 3 and 18 months. Moreover, such studies can now be designed to not just test for absolute differences between groups, but also to compare the rate of change over time with a repeated measures design, which we have established a baseline for here. Such studies promise to have more statistical power to detect small difference than designs that make discrete comparisons between 2 groups.

This study was also designed to test the hypothesis that visual aging is regulated by mitochondrial dysfunction. Our evidence that a drug (SS-31) that targets mitochondria and improves mitochondrial function is able to prevent and restore age related visual decline, is consistent that the hypothesis, and provides an incentive to determine how and where such a process may be occurring in the visual system. On the surface it may seem obvious to investigate whether mitochondrial changes in cones, or cone circuits in the retina would be the logical place to start. Indeed, it may be, but since visual behavior is a system level function, which involves retina, visual circuits in the brain downstream of the retina, and motor circuits that control smooth tracking responses, the ability to link a specific cell type or circuit with mitochondrial change that is manifest in visual behavioral change is challenge. It is possible that the effects we observed in our study are due to solely to changes in motor function, though we think that is not the most likely explanation. This is because the opto-kinetic tracking task we used here is based on titrating the salience of the visual stimulus (e.g. altering the spatial frequency or contrast), and then evaluating the motor response based on <u>any</u> evidence of (tracking). We are not in possession of technology to quantify the magnitude of the motor response during opto-kinetic tracking, which may reveal a role for a change in the motor response with age and treatment. Even if such a study revealed a reduction of motor system function with age, and an improvement with SS-31 treatment, it would not eliminate the role for a change in the ability of a visual sensory stimulus to alter opto-kinetic behavior in an age- and treatment-dependent way.

Exactly which visual circuits are involved in the changes reported are not known, but they are most likely resident in the retina and/or the accessory optic system in the brain (Sun et al, 2015), but not visual cortex. This is because previous studies have shown that the visual cortex is not part of the circuitry that normally enables opto-kinetic tracking. Our use of eye drops, in addition to s.c. injections, in this and a previous study (Alam et al, 2015a) represented a limited effort to direct the treatment to the eye and retina. That the effects of the eye drop treatments were comparable to those with s.c. injections in the study provides encouraging, but not conclusive evidence that the retina is involved in the changes. This is because it is possible that SS-31 delivered to the surface of the eye made its way to the rest of the body via the vascular tissues of the cornea and eyelid.

That we found photopic (cone) function to be selectively altered by age and treatment, however, provides an incentive to determine whether selective changes occur in retinal cones, or cone circuit function. Previous reports have provided evidence that SS-31 improves mitochondrial function by interacting with cardiolipin-rich membranes and enhancing formation of respiratory supercomplexes (Birk et al, 2013; 2014; 2015; Szeto, 2014). One strategy to link metabolic change to the effects we have reported here would therefore be to measure mitochondrial activity in the mouse retina and RPE-choroid-sclera complex, over the course of aging and with SS-31 treatment. Since cones comprise only ∼3% of mouse retinal photoreceptors (Jeon et al, 1998), however, it would be a challenge to dissociate rod contributions from cone contributions if metabolic changes were observed, or to definitively exclude a role for cones if no changes were found. In addition, that we found changes in temporal neural processing with age and SS-31 treatment in the study, may justify using an analysis of the flicker fusion electroretinogram (ERG) as an independent way to measure the temporal resolution capabilities of rod and cone photoreceptor systems (Dai et al, 2015). Indeed, when high frequency visual stimulation is used to produce an electroretinogram, rod responses are saturated by a flash and are not able to respond to a subsequent flash, and thus, the flicker analysis can be used to detect cone function. If such a flicker fusion study confirmed a selective ability of the cone system to change with age and SS-31 treatment, not only would it reinforce the findings of this study, but it would also link the changes to inner retinal circuits, which are through to enable flicker fusion function (Porciatti and Falsini, 1993).

### Clinical Relevance of SS-31 in Age-Related Ocular Disease

This study provides evidence that measures of spatial vision based on smooth tracking of moving stimuli can readily identify a treatable form of age-related visual decline. It also provides evidence that age-related decline of spatial visual function can be prevented and partially reversed. Since most measures of spatial visual function in humans don’t depend on smooth tracking-based assessments, but instead, rely on ‘stationary stimulus’ measures, such as identifying pictograms or letters on an acuity chart. Thus, it is not clear how our assessments and the effects of a therapy, would translate to humans. One way to bridge this gap would be to measure human age-related visual decline and the effects of a therapy using optokinetic procedures. We have recently introduced such a procedure (Mooney et al, 2018; 2020) which measures spatial vision based on smooth pursuit eye tracking. The application of this methodology to the measurement of age-related visual decline in humans should provide a more direct way to relate the preclinical findings in this study, to human studies. In the case of contrast sensitivity measures, which the methodologies are specialized to measure, they may be able to detect age-related visual dysfunction earlier in life, since contrast sensitivity is a more sensitive measure of spatial visual dysfunction than is acuity (Owsley, 2003). The method of drug administration is also of paramount consideration for any new patient intervention. Our results showed that eyedrops are a viable form of administration to treat age-related decline of temporal visual function. The recovery of photopic spatial acuity and contrast sensitivity after SS-31 eyedrops were comparable to daily s.c. injections. Eyedrops are probably a more tenable application approach both in the clinic, and for patient use in their home.

## Acknowledgements

We thank Stealth Biotherapeutics for providing SS-31, Hazel Szeto, Shaoyi Lui and Yi Soong, for laboratory support, and Peter Rabinovitch and James Hurley for their valuable insights on the project and manuscript.

